# X-ray Irradiation activates immune response in human T-lymphocytes by eliciting a Ca^2+^ signaling cascade

**DOI:** 10.1101/2020.11.13.379982

**Authors:** Dominique Tandl, Tim Sponagel, Sebastian Fuck, Timo Smit, Stephanie Hehlgans, Burkhard Jakob, Claudia Fournier, Franz Rödel, Bastian Roth, Anna Moroni, Gerhard Thiel

**Affiliations:** Department of Biology, Technische Universität Darmstadt, Darmstadt, Germany; Department of Radiotherapy and Oncology, Goethe-University, Frankfurt am Main, Germany; Department of Biophysics, GSI Helmholtzzentrum für Schwerionenforschung, Darmstadt, Germany; Department of Biosciences and CNR IBF-Mi, Università degli Studi di Milano, Milano, Italy

## Abstract

Radiation therapy is efficiently employed for eliminating cancer cells and reducing tumor growth. To further improving its therapeutic application it is mandatory to unravel the molecular effects of ionizing irradiation and to understand whether they support or counteract tumor therapy. Here we examine the impact of X-ray irradiation on immune activation of human T cells with single doses typically employed in tumor therapy. We discover that exposing cells to radiation triggers in a population of leukemic Jurkat T cells and in peripheral blood mononuclear cells (PBMCs) a canonical Ca^2+^ signaling cascade, which elicits immune activation of these cells. An early step in the signaling cascade is the initiation of sustained oscillations of the cytosolic Ca^2+^ concentration, an event mediated by store operated Ca^2+^ entry (SOCE) via an X-ray induced clustering of the Calcium Release-Activated Calcium Modulator 1 with the stromal interaction molecule 1 (Oari1/STIM1). A functional consequence of the Ca^2+^ signaling cascade is the translocation of the transcription factor nuclear factor of activated T cells (NFAT) from the cytosol into the nucleus where it elicits the expression of genes required for immune activation. These data imply that a direct activation of blood immune cells by ionizing irradiation has an impact on toxicity and therapeutic effects of radiation therapy.

## Introduction

Ionizing radiation (IR) is a universal tool in medical diagnostics and therapy. At higher doses (> 1 Gy) it serves as a major component in anti-tumor treatment (1). At lower doses (<1 Gy) radiationtherapy is mainly used for treatment of a variety of inflammatory/degenerative and hyperproliferative benign diseases (2, 3). A major dose limitation of radiation therapy is posed by toxic effects to the surrounding heathy tissue. In that context, peripheral blood mononuclear cells (PBMCs) cover an interesting population of cells. They are sensitive to IR (4,5) but, while cycling through the irradiated tissue, unavoidably exposed to it. Further, high dose exposure results in a suppression of immune functions (6). This includes among others killing of blood cells (7), induction of cell cycle arrests of immune cells (8) and triggering of pro-inflammatory processes (9, 10). By contrast, recent findings indicate an immune-stimulatory activity of high dose radiation with several studies describing a synergistic effect on local and distant tumor control when radiation therapy is combined with anti-checkpoint programmed death (PD-1) or its ligand PD-L1 immunotherapy (11,12). Finally, recent data suggest that irradiation at lower doses display anti-inflammatory or immune modulatory effects with consequences on immune surveillance of non-cancerous cells (13).

In a previous study we have reported that X-rays irradiation with low to medium dose elicits cellular responses, which are typically associated with immune stimulation in naïve T-lymphocytes (14). These include an increase in cell diameter, up-regulation of CD25 membrane expression, interleukin-2 and interferon-gamma synthesis, an enhanced integrin-mediated adhesion of Jurkat cells, as model for T cells, or PBMCs to endothelial cells. Since many of these mechanisms are mediated by a Ca^2+^ signaling cascade we anticipate a causal interrelationship between radiation stress and induction of a Ca^2+^ signaling cascade (5, 14). Here we aimed to examine whether clinically relevant doses of X-ray between 0.5 and 5 Gy trigger Ca^2+^ signaling events in T cells and whether these signal transduction cascades are relevant for immune stimulation. We observed that X-ray doses <1 Gy triggered in Jurkat cells long lasting episodes of Ca^2+^ oscillations with a delay of some 10 minutes. These oscillations are mediated by *store operated Ca*^2+^ *entry* (SOCE) via a radiation-induced clustering of Calcium Release-Activated Calcium Modulator 1 and the stromal interaction molecule 1 (Orai1/STIM1) in the plasma membrane. Like immune stimulation by antigens also X-ray exposure induced Ca^2+^ entry via the SOCE pathway, which in turn stimulates the nuclear translocation of the transcription factor *nuclear factor of activated T cells* (NFAT). This stimulus-induced and Ca^2+^-dependent nuclear import is a well-known key step for cytokine production, proliferation and immune competence (15).

## Results

### Ionizing irradiation elicits a delayed Ca^2+^ response in Jurkat cells

It was previously shown that X-ray irradiation triggers an immune stimulation in T cells, which is mediated by an increase in the concentration of free Ca^2+^ in the cytoplasm (Ca^2+^_cyt_) (14). To unravel a causal relationship between IR and Ca^2+^_cyt_ signaling cascades we monitored the immediate impact of X-ray exposure on the level of this second messenger in Jurkat cells. Cells loaded with the fluorescent Ca^2+^ reporter dye Fluo-4 were imaged in real-time on a fluorescence microscope directly coupled to an X-ray source. Fig. 1A shows that X-ray doses of 1 and 10 Gy did not elevate Ca^2+^_cyt_ levels within 10 min post irradiation. Combined with the finding that Ca^2+^_cyt_ in the same cells can be elevated by ionomycin (Fig. 1A) the data suggest that IR exposure has no immediate impact on Ca^2+^_cyt_ concentrations in Jurkat cells.

**Figure 1.**
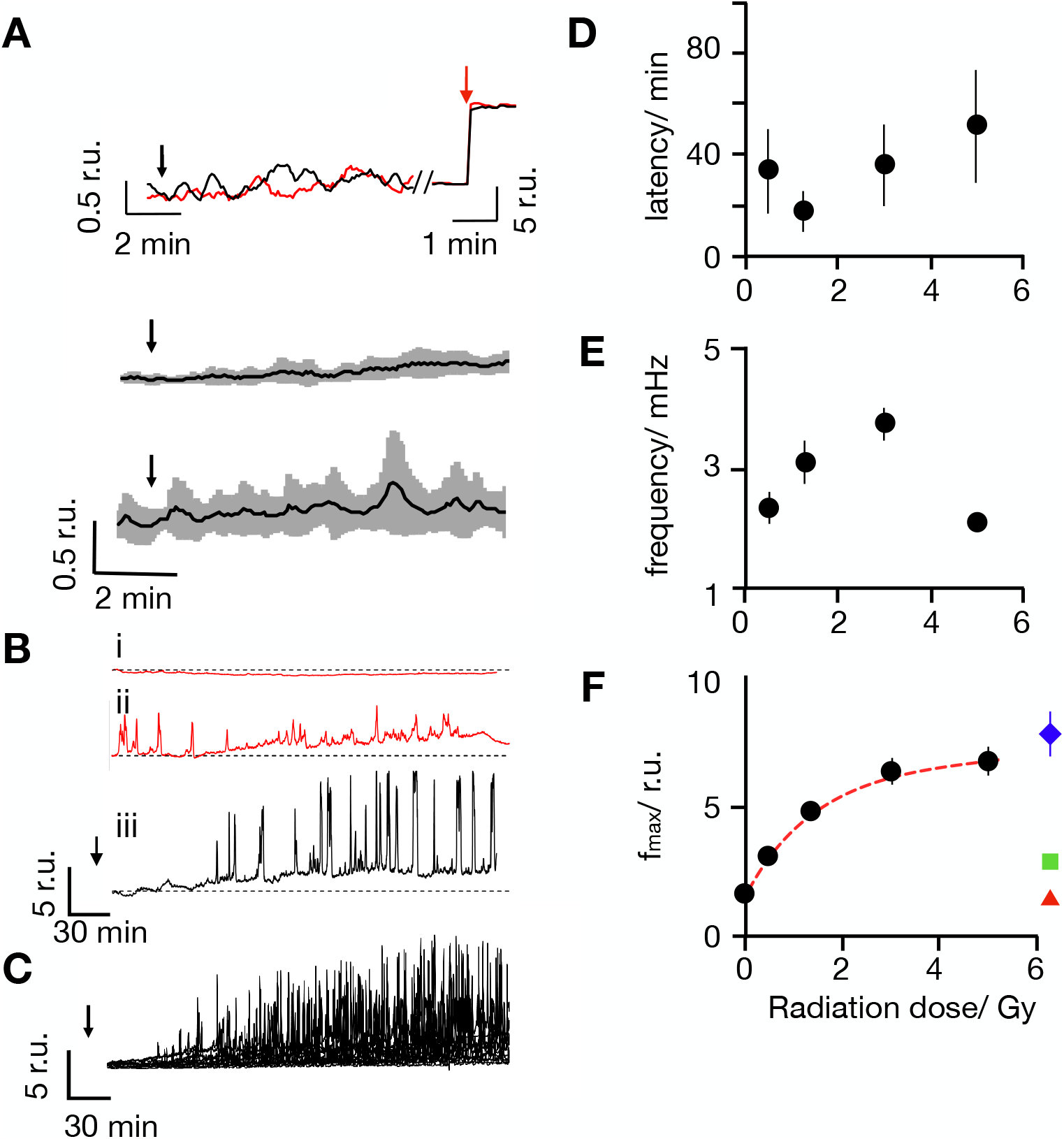
Ionizing irradiation elicits delayed Ca^2+^_cyt_ oscillations with distinct frequencies and amplitudes. (A) Representative Fluo-4 signals as measured as changes in cytosolic Ca^2+^_cyt_ in Jurkat cells (Top). Fluorescence was recorded in real-time before and after irradiation with 1 Gy (red) and 10 Gy (black) at times indicated by black arrows. The same cells responded with a maximal fluorescence increase after addition of 1 μM ionomycin (red arrow). Mean Fluo-4 intensity from cells irradiated at arrow with 1 Gy(middle) or 10 Gy (bottom) X-ray. Data are means ± SD (grey) from 15 cells for each dose. (B) Representative long-term measurements of Fluo-4 intensity in Jurkat cells of which two were non irradiated (i. ii, control, red) and the other exposed to a dose of 5 Gy X-ray (iii, black). Time of X-ray exposure is indicated by black arrows. While control cells maintain a stable Fluo-4 signal (i) or reveal irregular oscillations (ii) the irradiated cell starts to oscillate after a lag time (iii). (C) Overlay of Fluo-4 signal from 15 individual cells after irradiation with 5 Gy; Time of X-ray exposure is indicated by black arrows.Latency time between onset of Ca^2+^_cyt_ oscillations after irradiation (D) oscillation frequency (E) and maximal amplitude of oscillation as a function of irradiation dose (F). Data are mean values ± SD from >25 cells per dose. All Ca^2+^_cyt_ measurements in response to X-irradiation were performed in buffer containing 2 mM Ca^2+^. The colored symbols show the maximal level of Fluo-4 fluorescence intensity obtained in the same buffer by depleting Ca^2+^ stores with 2 μM thapsigargin (blue triangle) or by adding 1 μM ionomycin (green square); red triangles report Fluo-4 intensity elicited by 5 Gy in Ca^2+^ free external buffer.

To capture potentially delayed Ca^2+^_cyt_ responses Fluo-4 fluorescence was monitored over an extended time window. Representative recordings in Fig. 1B indicate that most untreated cells maintained a constant low Ca^2+^_cyt_ over some hours of recording (Fig 1Bi). In 130 control cells we only observed in 20% of the cells some spontaneous and non-periodic excursions in Ca^2+^_cyt_; the latter were mostly visible already at the start of the imaging (Fig. 1Bii). After exposure to 5 Gy approximately 50% of the cells exhibited characteristic delayed Ca^2+^_cyt_ oscillations. In the representative example in Fig.1Biii the cell started oscillating after a lag time of 65 min (Fig. 1B). Similar delayed and long lasting Ca^2+^_cyt_ oscillations were observed in a larger fraction of cells after irradiation with 5 Gy (Fig. 1C). The remaining cells either maintained a constant Ca^2+^_cyt_ or exhibited unspecific Ca^2+^_cyt_ excursions as in control cells (Fig. 1Bi,ii). For quantifying the probability of radiation-induced effects we consider here only Ca^2+^ oscillations, which occurred ≥10 min after start of the imaging. From this analysis it occurs that oscillations are only visible in ca. 10% of the control cells but in 56% of cells after irradiation with a dose of 5 Gy (Fig. S1A).

Experiments were repeated over a range of X-ray doses from 0.5 to 5 Gy. These treatments also elicited with a high probability delayed Ca^2+^_cyt_ oscillations (Fig. S1A). The lag time between X-ray stimulation and onset of Ca^2+^_cyt_ signaling events varied considerable from one cell to the other. In 70% of the stimulated cells with 1 Gy the first detectable Ca^2+^ peak occurred between 10 min (fastest) and 72 min (slowest) after X-ray exposure. A plot of the average lag times as a function of X-ray dose indicates that this value is not significantly changing with the stimulation dose (Fig. 1D). A frequency analysis further reveals that Ca^2+^_cyt_ oscillates in response to X-ray with 2 to 4 mHz. (Fig. 1E). This frequency remains largely constant over the range of irradiation doses.

The traces depicted in Fig. 1Biii, C show that the amplitude of the Ca^2+^_cyt_ excursions increases gradually with a saturation kinetics after onset of the oscillations. The peak values of the maximal Ca^2+^_cyt_ excursions are a function of the stimulation doses. From a fit of the data with a single saturating exponential function:

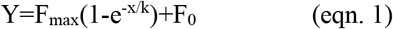

where F_max_ is the maximal increase in peak fluorescence, F_0_ the background fluorescence of the control and k the dose for half maximal increase in fluorescence the half maximal Ca^2+^_cyt_ peak is achieved by 1.5 Gy (Fig. 1F).

In the case of antigen-mediated T cell activation, Ca^2+^_cyt_ oscillations are the result of *store operated Ca*^2+^ *entry* (SOCE), a mechanism, which can be imitated by thapsigargin (TG), an inhibitor for sarcoplasmic/endoplasmic reticulum calcium ATPase (SERCA). In Jurkat cells treated with 2 μM TG the Ca^2+^_cyt_ signal increases to a new plateau; the latter, however, remains well below that of the maximal amplitude measured in response to X-ray stimulation. Notably, the maximal amplitude value was in the same range of those elicited by the Ca^2+^ ionophore ionomycin (Fig. 1F). This indicates that IR elicits Ca^2+^_cyt_ excursions to much higher levels than TG; the former amplitudes presumably exceeded the saturation of the Fluo-4 dye, which is about 1 μM (16, 17).

### Ca^2+^_cyt_ oscillations can be suppressed by buffering external Ca^2+^ and by blocking Ca^2+^ influx

Stimulus-induced Ca^2+^_cyt_ oscillations can originate from a release of Ca^2+^ from internal stores or from entry via a plethora of Ca^2+^ permeable channels in the plasma membrane of T cells (18). To test the contribution of plasma membrane channels to this process, experiments similar to those in Fig. 1 were repeated in a nominally Ca^2+^ free extracellular solution. In these experiments the probability of finding Ca^2+^_cyt_ oscillations was reduced to that of untreated control cells by the addition of 5 mM EGTA to the buffer (Fig.S1A). We further measured the mean fluorescence over a time window of 60 min to 120 min after X-ray exposure. The value was significantly higher in cells treated with 5 Gy X-ray than that of the sham-irradiated control group (Fig. S1B). In the presence of EGTA this value was greatly reduced and only 2 times higher than the control. The results of these experiments show that the absence of external Ca^2+^ abolishes radiation induced Ca^2+^_cyt_ oscillations but still allows some steady increase in Ca^2+^_cyt_. This suggests a calcium influx via plasma membrane channels as the main trigger of Ca^2+^_cyt_ oscillations in irradiated cells. To further test this hypothesis, experiments were repeated in a buffer containing Ca^2+^ and ± gadolinium (Gd^3+^), a broad inhibitor of Ca^2+^ permeable channels in T cells (19,20). In the presence of 5 μM inhibitor the probability of radiation-induced Ca^2+^_cyt_ oscillations was reduced close to that in control cells (Fig. 1SA). Like EGTA Gd^3+^ did not fully prevent a radiation induced increase in the concentration of Ca^2+^_cyt_ above the sham-irradiated control (Fig. S1B). Together these data underscore the impact of radiation on Ca^2+^_cyt_ and the importance of Ca^2+^ influx for the radiation-induced signaling cascade, which leads to Ca^2+^_cyt_ oscillations.

### Ca^2+^ oscillations are mediated by STIM/Orai activation

The major mechanism for Ca^2+^ entry into T cells is provided by the *calcium induced calcium release* (CICR) pathway (18). In this system the calcium level of endoplasmatic reticulum (ER) stores is monitored by the ER Ca^2+^ sensors stromal interaction molecules (STIM). Upon store depletion STIMs aggregate and move to contact points with the plasma membrane (PM) where they interact and activate Orai channels (21). The active Orai1/STIM1 complex, which assembles to the CRAC channel (22), is necessary and sufficient to support SOCE. To test the involvement of this pathway in the radiation induced T cell stimulation we monitored the dynamic distribution of Orai1 and STIM1 in Jurkat cells and PBMCs. Cells were fixed for immunostaining 15, 30, 60 min after X-ray exposure or activating with TG. The representative images in Fig. 2A and Fig. S2 show the typical distribution of the two components in unstimulated cells. The Orai channel subunit is evenly distributed in the PM while the STIM protein generates a diffuse signal throughout the cell (Fig. 2, Fig. S2). After activating cells with 2 μM TG, STIM proteins are translocated from the cytosol to the plasma membrane where they colocalize with the Orai proteins in distinct clusters.

**Figure 2.**
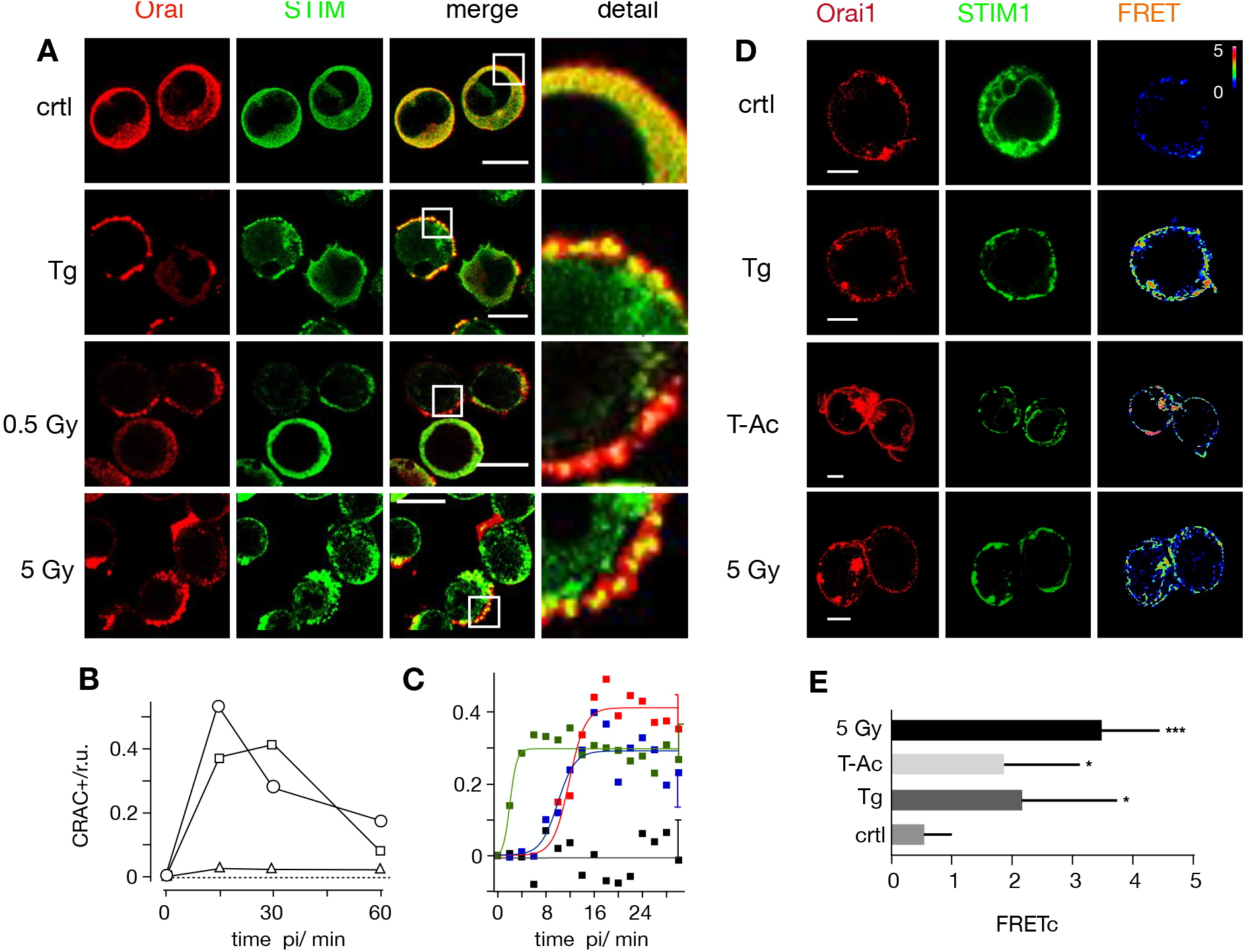
Ionizing radiation triggers Ca^2+^ regulated STIM1/Orai1 CRAC channel formation. Distribution of endogenous STIM1 (green 1^st^ column) and Orai1 (red 2^nd^ column) in Jurkat cells immunostained with Alexa (Alx)488 and Alx647, respectively. An overlay of green and red channels with magnification of indicated area are shown in 3^rd^ and 4^th^ columns. Fixed cells were obtained from untreated cells (top row), cell treated for 15 min with 2 μM thapsigargin (2^nd^ row) or from cells 15 min after X-ray exposure to 0.5 Gy (third row) or 5 Gy (bottom row). (B) Frequency of Jurkat cells with positive clustering of STIM/Orai1 after store-depletion by irradiation with 0.5 Gy X-ray (triangles), 1.5 Gy (squares) or 5 Gy (circles). For each condition ≥ 282 cells were analyzed. (C) Pearson correlation coefficient (PC) for colocalization of STIM1-eYFP and Orai1-eCFP. Data obtained from confocal live cell real-time acquisition of Jurkat cells heterologously expressing the two proteins. After normalizing the start value of all treatments, the PC value remains at the same level in control cells (black) but increases with different kinetics in cells stimulated with 2 μM thapsigargin (green), 25 μL/mL activator (blue) or 5 Gy (red) X-rays. The data were fitted with logistic equation (eqn 1, solid lines) yielding the following times for maximal increase in STIM1/Orai colocalization: 2 min TG, 10 min T-Ac, 12 min X-ray. Data are mean values ±SD from ≥ 5 independent experiments. The mean SD of all data points is shown on the last data points. All scale bars10 μm. (D) Representative confocal images of same cells with fluorescent donor molecule Orai1-eCFP (red, 1^st^ column), acceptor molecule STIM1-eYFP (green, 2^nd^ column) and heat maps of the resulting FRET signals (3^rd^ column). Images are from untreated cells (control), or cells incubated with 2 μM thapsigargin, 25 μL/mL activator (T-Ac) or irradiated with 5 Gy. Cells were fixed 15 min after start of treatment. All three treatments generate a visible FRET-signal in the plasma membrane. Scale bars, 10 μm. (E) Mean FRET signal (± SD, n≥5 cells) from plasma membrane of cells treated as in D: control cells (crtl), 5 min in 2 μM Tg, 15 min in 25μl/ml T-Ac or 20 min post irradiation with 5 Gy. The FRET signal from treated cells is significantly higher than that of the control value (*P < 0.05, ***P < 0.001 from Student’s-test)

The same distinct overlay of STIM and Orai in plasma membrane puncta emerged after exposing Jurkat cells to a dose of 5 Gy (Fig. 2A). Already 15 minutes after irradiation a maximum clustering of both proteins was observed in approximetly 50% of the cells analyzed before the signal gradually decreased (Fig. 2B). A similar transient clustering of STIM and Orai was evident with a 1.5 Gy X-ray exposure. In experiments with 0.5 Gy it was still possible to detect individual cells with a clear clustering of the two proteins (Fig. 2B, C). This confirms that low dose irradiation is able to elicit STIM/Orai aggregation; the numbers however were too low for a robust statistical analysis.

To confirm X-ray triggered aggregation of STIM and Orai we co-expressed STIM1::eYFP and Oari1::eCFP in Jurkat cells and measured protein/protein interactions by Förster resonance energy transfer (FRET). Data in Figs 2D,E show that the FRET signal is small in untreated control cells. It significantly increased in cells stimulated with 2 μM TG or 25 μL/mL ImmunoCult human CD3/CD28/CD2 T cell activator (T-Ac). An even larger energy transfer was measured in cells exposed to a dose of 5 Gy. Together these data confirm that STIM1 and Orai1 interact in response to X-ray exposure.

To estimate the time course for stimulus-induced STIM1/Orai1 clustering we measured in cells, which co-express both proteins, the Pearson correlation coefficient (PCC) for the yellow fluorescent STIM1 and the cyan fluorescent Orai1 after stimulation. The mean colocalization values in unstimulated cells were low (PCC=0.3 ± 0.08) and did not change over time (Fig. 2C). After stimulation the PCC value increased rapidly in response to TG (PCC=0.76 ± 0.05) and slow after adding the activator T-Ac (PCC=0.79 ± 0.04). The irradiation-induced increase in STIM1/Orai1 cluster formation progressed with the same slow kinetics of the activator to a maximal PCC value of 0.79 ± 0.06. The kinetics of changes in the PCC values could be fitted by a logistic equation:

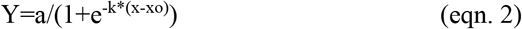

where a is the maximal PCC value, k the rate of increase and x_o_ the time of maximal increase.

In case of the 5 Gy exposure the maximal increase in co-localization was achieved ~12 min after stimulation (Fig. 2C). This value is in good agreement with the mean delay between X-ray stimulation and the onset of Ca^2+^_cyt_ oscillations as depicted in Fig. 1.

### The STIM/Orai activation pathway is also activated in naϊve T-lymphocytes

Jurkat cells are a leukemic T cell line, which serves as a model system for uncovering the basic signaling events engaged in T cell activation (23). To test whether the irradiation triggered Ca^2+^ signaling cascade is also occurring in naïve T cells, we repeated the experiments in Fig. 2 with PBMCs. The images in Fig. 3A indicate that the cytosol volume of non-stimulated lymphocytes is even smaller than that of Jurkat cells, which makes it more difficult to detect a translocation of STIM from the cytosol to the membrane resident Orai. A comparative analysis nonetheless shows that the Orai and STIM distribution remains uniform in unstimulated control cells while they exhibit a distinct clustering after stimulation with 2 μM TG. The same clustering is also evident after exposing cells to 5 Gy (Fig. 3A). This underscores that the IR induced Ca^2+^ signaling cascade is a general response of resting T cells.

**Figure 3.**
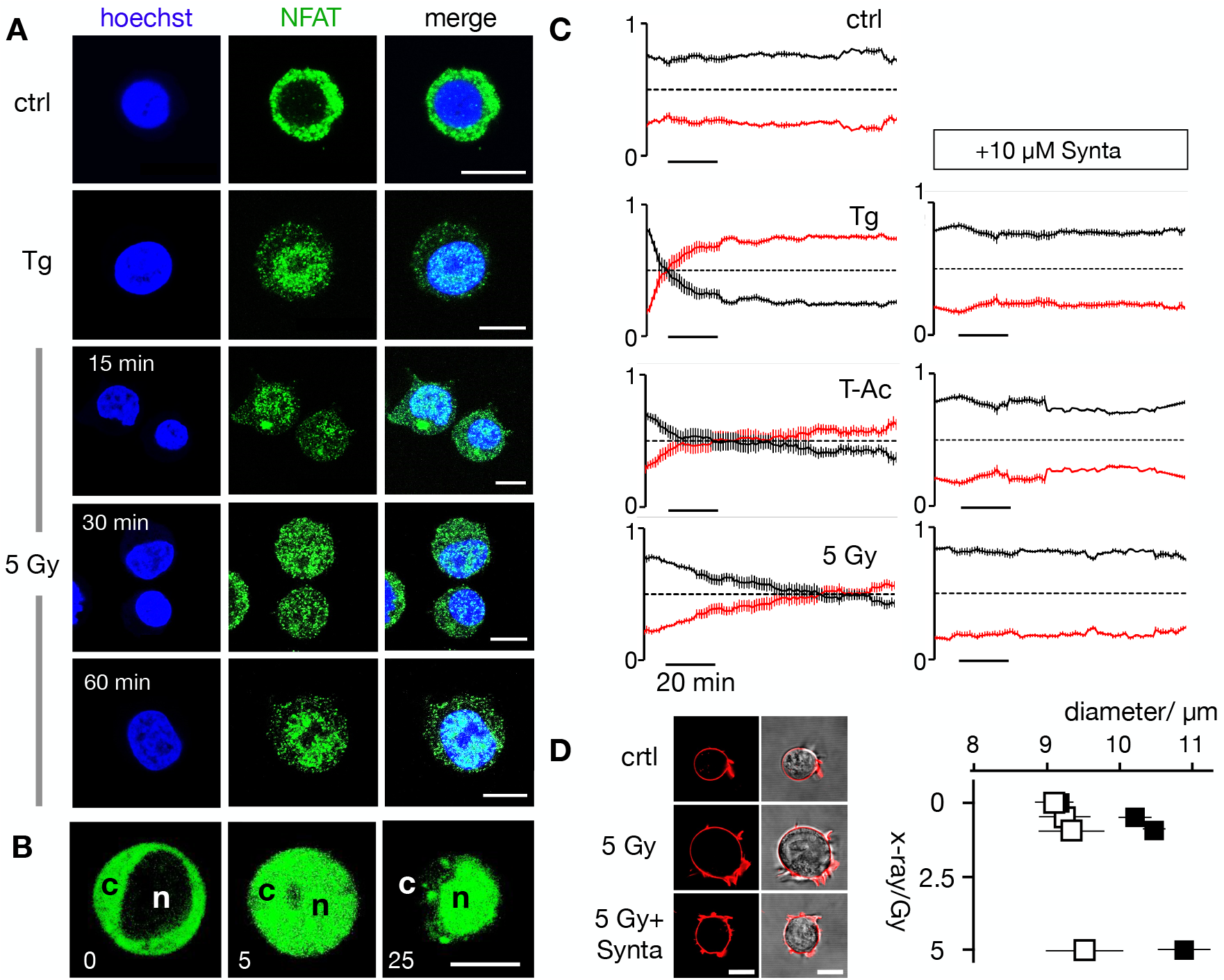
Calcium-dependent SOCE/NFAT pathway is activated by IR in naϊve T-lymphocytes. (A) Distribution auf endogenous STIM1 (green, left column) and Orai1 (red, 2^nd^ column) in fixed PBMCs immune-stained with Alx488 and Alx647 respectively. A merger of the two channels is shown in 3^rd^ column with a higher magnification of the marked area in the right column. In control cells also the nucleus is additionally stained with Hoechst dye. Cells were fixed (control cells, top) 15 min after treatment with 2 μM thapsigargin (Top) or 15 min after exposing cells to either doses of 0.5 Gy or 5 Gy. (B) Confocal images of endogenous NFATc2 (green) stained with Alx488 (left column) and Hoechst DNA dye (2^nd^) in PBMC. Overlay of both columns is shown in the third row. Cells were fixed immediately (control, top), 15 min after 2 μM thapsigargin Ca^2+^ store depletion (middle) or 60 min after X-ray exposure with 5 Gy (Bottom). All scale bars,10 μm.

### Ca^2+^ signaling cascade results in a translocation of NFAT to the nucleus

Ca^2+^ entry via the SOCE pathway is the main source for activating the transcription factor isoforms of NFAT (24, 25). The stimulus-induced and Ca^2+^-dependent nuclear translocation of NFAT is instrumental for subsequent cytokine expression, proliferation and immune competence (26). Notably, the frequency of the X-ray induced Ca^2+^ oscillations in Jurkat cells (Fig. 1E) is typical for a signaling cascade, which elicits the activation of the NFAT pathway (27).

To test if the NFATc2 pathway is indeed activated by ionizing radiation we monitored nuclear translocation of endogenous NFAT labelled with Alexa (Alx)488. The representative images depicted in Fig. 3B and 4A show that the transcription factor is primarily located in the cytosol in unstimulated Jurkat cells (4A) and naϊve T cells (3B). Stimulation with 2 μM TG favors a translocation of NFAT into the nucleus in a population of cells tested (Fig. S1C). The similar nuclear translocation is induced in 72% of cells irradiated with 5 Gy(Fig. 4A,B, Fig. S1C).

**Figure 4.**
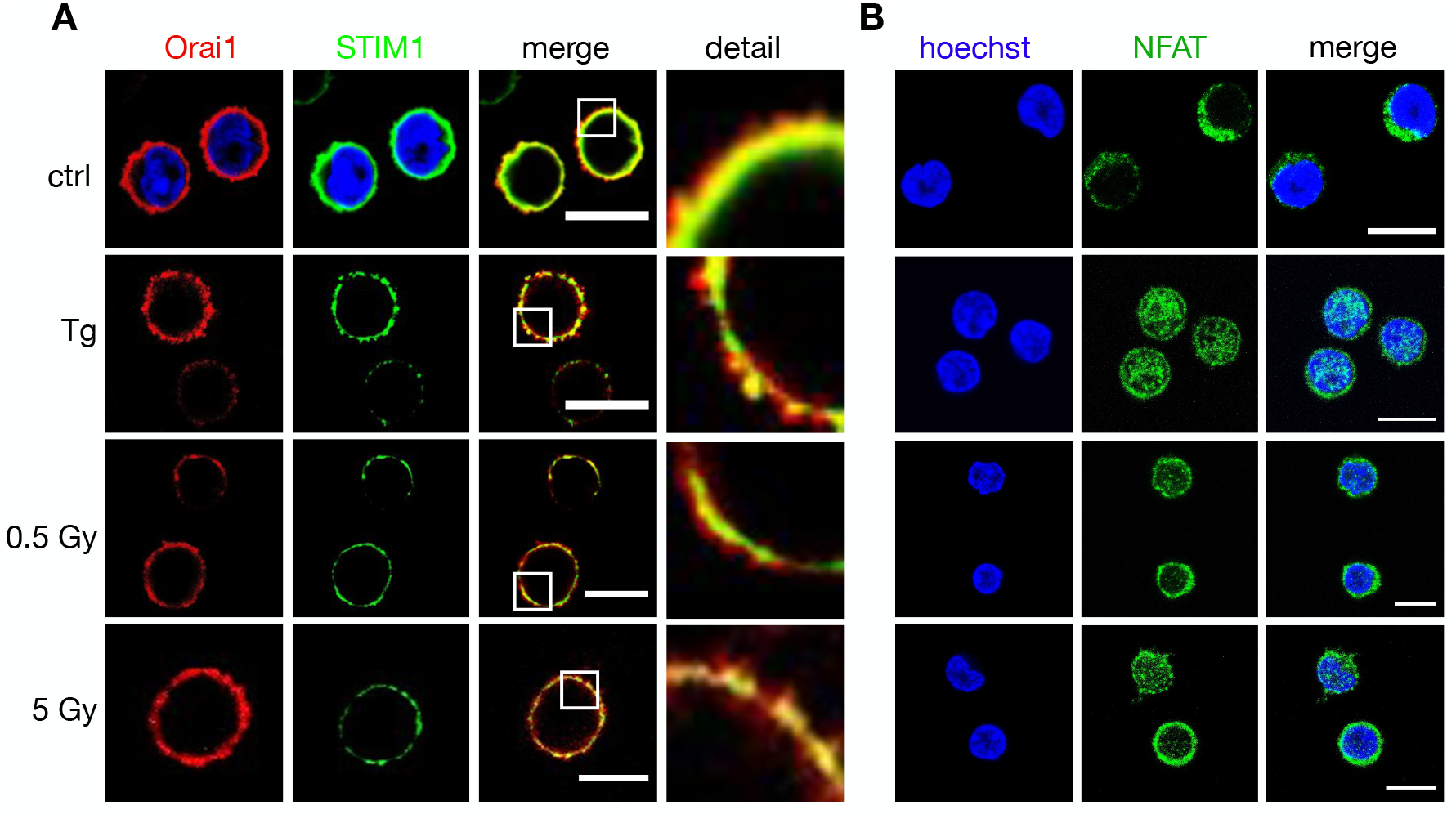
Nuclear translocation of Ca^2+^-dependent NFAT in Jurkat cells. (A) Confocal images of endogenous NFATc2 (green) immunostained with Alx488 (green, left row) and Hoechst DNA dye (blue, 2^nd^ row) in Jurkat cells. The third row shows a merge of blue and green channels. Cells were fixed immediately in control cells (Top), 15 min after 2 μM thapsigargin Ca^2+^ store depletion (second panel) or 30, 60 min after X-ray exposure with 5 Gy (two bottom panels). (B) Life cell imaging of nuclear import of transiently expressed NFATc2-GFP from cytosol (c) to nucleus (n) in Jurkat cells after stimulation with 2 μM thapsigargin. (C) Kinetic analysis of NFATc2-GFP nuclear import reactions (nucleus red, cytosol black) from confocal imaging of Jurkat cells in control condition (crtl), in 2 μM thapsigargin (Tg) in 25 μL/mL activator (T-Ac), or after a 5 Gy exposure (5 Gy). Data were obtained without (left) and with 10 μM CRAC channel inhibitor Synta (right). Each time course diagram is the mean ± S.E. of ≥ 12 individually measured cells. (D) Jurkat cells exhibit a dose-dependent increase in diameter 24 h after irradiation, which can be abolished by 10 μM Synta. Representative images (left, confocal image of fluorescent stained plasma membrane (red) and overlay of fluorescent and bright field image) of individual Jurkat cells. Images were taken 24 after start of experiment with an untreated cell (ctrl), a cell exposed to Gy X-ray without (5Gy) and with 10 μM Synta (5Gy+Synta). Mean diameters ± SD of > 300 cells per treatment 24 h after exposed to X-ray between 0 and 5Gy in the absence (black) or presence (open) 10 μM Synta. The diameters ± Synta are significantly different (*P < 0.05, **P < 0.01, ***P < 0.001 from student t-test). All scale bars,10 μm.

Stimulus-induced translocation of the heterologous expressed GFP tagged NFAT can be monitored with live cell imaging by following its translocation dynamics in real-time (Fig. 4B). Again, analyses of NFAT accumulation in the nucleus highlights distinct response times to different stimuli. While TG stimulation causes a 50% nuclear translocation of NFAT (in 97% off all cells monitored) after 8 ± 1.5 min, it takes more than 21 ± 6.5 minutes for a comparable response in 89% of the cells stimulated with T-Ac and 70 ± 5 minutes for a 5 Gy irradiation 67% of all cells (Fig. 4C). The nuclear translocation of NFAT is causally related to the activation of CRAC channels for all three stimuli as treating cells with the CRAC channel inhibitor Synta abolishes the nuclear translocation of NFAT (Fig. 4C; Fig. S1C) in all cases.

Finally, to examine the consequence of CRAC channel activation on the physiological response of Jurkat cells we examined the effect of 10 μM Synta on irradiation induced increase in cell diameter; since the morphological change is part of the radiation-induced immune response (14). The data depicted in Fig. 4D indicate that the cells have a uniform diameter under control conditions. Over 24 h, this diameter is not changing neither in the presence of 10 μM Synta nor in the absence of Synta. X-ray irradiation with three different doses (0.5, 1.5 and 5 Gy) caused a dose-dependent increase in cell diameter. This increase in cell size was largely abolished in the presence of the CRAC channel blocker (Fig. 3D). The results of these experiments suggest that the morphological change in Jurkat cells in response to X-ray stimulation is an endpoint of a signal transduction cascade, which involves SOCE via STIM1/Orai1 clustering with subsequent NFATc2 activation.

## Discussion

Stimulus induced Ca^2+^ signaling cascades are a key event in the activation of T cells. Triggered by antigen binding to the T cell receptor (28), a downstream signaling cascade is initiated, which promotes the release of Ca^2+^ from internal stores and eventually the activation of SOCE. In this process Ca^2+^ enters the cytoplasm primarily via CRAC channels and generates Ca^2+^_cyt_ oscillations with distinct durations and amplitudes (29,30). These dynamic changes in the concentration of the second messenger molecule are finally decoded by cytosolic Ca^2+^-dependent target enzymes including kinases, phosphatases and transcription factors such a NFAT (31-33). With this network of signaling steps T cells achieve precise control over essential lymphocyte functions such as cytokine production, proliferation, differentiation and antigen dependent cytotoxicity. In the present study, we indicate that the signaling cascades involving CRAC channel activation, SOCE-mediated Ca^2+^_cyt_ excursions and translocation of NFAT from the cytosol to the nucleus can be triggered in a population of T cells by clinically relevant doses of ionizing irradiation. In this regard IR elicts comparable effects to a well-established activator of T cells or of TG. Further, the finding that a specific inhibitor of CRAC channels blocks the crucial translocation of NFAT and physiological responses of T cells irrespectively of the nature of stimulation underpins that they all employ SOCE as the major Ca^2+^ entry pathway. Other Ca^2+^ channels in T cells seem to play no primary role in IR-induced Ca^2+^ signaling.

It is well established that the form of Ca^2+^_cyt_ signals in T cells determines their cellular response (30, 34-36). Depending on the nature and the concentration of the stimuli the Ca^2+^ signal can either exhibit a sustained increase in Ca^2+^_cyt_ or periods of Ca^2+^_cyt_-oscillations with different frequencies and amplitudes. This frequency and amplitude encoded signature of the Ca^2+^_cyt_ oscillations bears information on the subsequent differentiation of T cells; it can induce proliferation or cell death (37). Different amplitudes and frequencies also activate different transcription programs in populations of T cells (38). Scrutiny of IR-nduced Ca^2+^ oscillations show that they oscillate independent on the IR doses with frequencies of 2-4 mHz. This is in the frequency range of antigen-triggered oscillations in T cells which eventually result in an activation of NFAT (24, 27). Our finding that IR activates the NFAT pathway in T cells is in good agreement with the view that both stimuli trigger the same signaling pathways.

A peculiarity of IR triggered Ca^2+^_cyt_ oscillations is their high amplitude. They reach the same level as those induced by ionomycin suggesting that the peak amplitudes must exceed Ca^2+^_cyt_ concentrations of 1 μM e.g. the concentration at which the fluorescent sensor approaches saturation. The amplitudes of TG-induced excursions of Ca^2+^_cyt_ are well below this value suggesting that the response of T cells to IR is similar but not identical to a canonical antigen stimulation by T-Cell activator. This difference in the Ca^2+^ code may contribute to the findings that IR does not elicit the same degree of immune activation in T cells than the activator (14). This might bear consequences for the spectrum of physiological reactions, which are triggered by IR. Important to consider in this context is that a high Ca^2+^ load can induce cell death in lymphocytes. Together with the finding that the amplitude of the Ca^2+^_cyt_ oscillations increase with the IR, this might be a possible explanation for an induced death of T cells by high dose IR exposure (39).

An interesting finding in the present study is that IR induced Ca^2+^_cyt_ oscillations are not an immediate consequence of radiation exposure; they only occur with a delay of several minutes. This delay is well beyond the life-time of oxygen radicals including the long lived H_2_O_2_ (40, 41) meaning that Ca^2+^ oscillations are not initiated by radiolysis of water or peroxidation/oxidation of lipids/proteins. This lack of an immediate impact of IR on Ca^2+^_cyt_ is consistent with previous high-resolution imaging experiments which showed that individual high energy ions had no immediate impact on the Ca^2+^ concentration along the track of the ions (42). The current data are not sufficient to explain the gap between IR exposure and the onset of Ca^2+^_cyt_ oscillations. Worth noting, however, is that typical T cell stimuli like phytohemagglutinin are not causing an immediate Ca^2+^ response (43); like in the case of IR it takes up to 5 min before Ca^2+^_cyt_ starts to oscillate. This parallel in the dynamics between antigen and IR-induced events is even more evident in the time course of STIM/Orai clustering, which shows about the same time course irrespectively of a stimulation by IR or by a T cell activator. If these parallels are a coincidence or based on a common mechanism like generation of inositoltrisphosphate (IP_3_) and/or diazylglycerol (DAG) upstream of SOCE needs to be investigated.

Since immune cells like T cells, which are cycling in the blood, are inevitably exposed to IR during tumor therapy, it is important to understand their response to IR. The present study underpins a stimulating role of IR on STIM/Orai cluster formation and a subsequent activation of CRAC channels for SOCE not only in a T cell cancer cell line (Jurkat cells), but also in naïve peripheral blood lymphocytes. T-lymphocyte activation is in the forefront of anti-tumor cytotoxic effects and the regulation of an adaptive immune response in healthy and cancerous tissue (44,45). Our findings thus have implications for the understanding of both, radiation associated toxicity in normal tissue and the efficacy of radiation therapy, especially if combined with checkpoint PD1 and PD-L1 inhibitors in current clinical practice. Further, STIM and Orai proteins are not restricted to T cells but also expressed in B-cells as well as in the phagocytes such as neutrophilic granulocytes, macrophages, and dendritic cells (46) where they regulate a multitude of cellular reactions (15). With a more general functional importance of these channel forming proteins in different types of immune cells we anticipate that IR activation of CRAC channels will even have a more global importance in the modulation of the immune responses following IR.

## Materials and Methods

### Cell culture

Jurkat cells (ACC 282) were purchased from the German Collection of Microorganisms and Cell Cultures (DSMZ, Braunschweig, Germany). They were grown in RPMI 1640 medium (Thermo Fischer Scientific Waltham, MA, USA), supplemented with 10% heat inactivated fetal calf serum (FCS; PAA, Cölbe, Germany) and 50 U/ml penicillin plus 5 μg/ml streptomycin (Sigma-Aldrich, Munich, Germany). Peripheral blood mononuclear cell (PBMCs) were isolated from blood of healthy volunteers, using density-gradient centrifugation (Biocoll Separating Solution, Biochrom, Berlin, Germany) and maintained in RPMI 1640 Medium with 10% FCS, plus 50 U/ml penicillin and 5 μg/ml streptomycin prior to assays as described previously (14).

### Cell irradiation and treatments

Cells were exposed to X-ray irradiation in T_35_ petri dishes using an Isovolt 160 Titan E source with a voltage of 90 kV and 33.7 mA (GE Sensing & Inspection Technologies, Alzenau, Germany), with a dose rate of 0.055 Gy/s. Ionomycin (# ab120370, Abcam, Cambridge, UK), thapsigargin (TG) (Sigma-Aldrich, Taufkirchen, Germany) and the cell-permeable Ca^2+^ sensor Fluo-4 AM (Life Technologies, Carlsbad, CA, USA) were dissolved in DMSO and immediately added to external solution prior to experiments in a final concentration of 5/2/1 μM, respectively. To activate human T cells, ImmunoCult™ Human CD3/CD28/CD2 T cell activator (short T-Ac) (Stem cell Technologies, Vancouver, BC, Canada) was added to the cell culture medium (25 μL per 1 mL of cell suspension). The cell permeable organelle tracker Mito-Tracker Green FM (#M7514) and ER-Tracker Red (#E34250) and PM tracker CellMaskOrange (Thermo Fisher Scientifc) were used according to the manufacturer’s recommendations. Cell nuclei were stained with Hoechst dye (5 μg/ml, Thermo Fisher Scientifc) diluted in external microscopy buffer, for 10 min at 37°C. Subsequently, cells were washed and resuspended in dye-free microscopy buffer.

### Determination of cell diameters

Jurkat cell diameters were measured with an EVE automatic cell counter (NanoEnTek, Seoul, South Korea) and corrected manually using EVE PC.LNK 1.0.3 software. Cell viability was determined by using trypan blue exclusion.

### Confocal laser scanning microscopy

Confocal laser scanning microscopy was performed on a Leica TCS SP or SP5 II system (Leica microsystems, Mannheim, Germany) equipped with a 40 × 1.30 oil UV (HCX PL APO), a 63 × 1.4 oil UV (HCX PL APO lambda blue) or a 100 × 1.44 oil UV objective (HCX PL APO CS). The external buffer used for microscopy (MB) contained in mM: 140 NaCl, 4 KCl, 1 MgCl_2_, 5 Mannitol, 10 HEPES, 2 CaCl_2_, pH 7.4 with osmolarity of 310 mosmol/L. Live cell imaging of changes in Ca^2+^_cyt_ and translocation of NFAT-GFP were performed as described in (14) and (47) respectively. STIM1-eYFP and ORAI-eCFP were transiently expressed as described in (21). To enable a gentle adhesion of the cells to the glass coverslips (Ø 25 mm) they were prepared by cleaning in a plasma furnace (Zepto-B, Diener electronic GmbH, Ebhausen, Germany) and coated with one layer of PBS/5% BSA in a spincoater (PIN150, SPS Europe Spincoating, Putten, Netherlands) and a second layer of 0.01% poly-L-lysine (molecular weight 75–150 kDa).

For monitoring the Ca^2+^_cyt_ Jurkats were loaded with the cell permeable Ca^2+^ sensor Fluo-4 (Life Technologies) for 30 min in microscopy buffer in a final concentration of 1 μM. The calcium dye was subsequently removed by washing cells with dye-free buffer. Calcium signals were recorded in a time interval of 5 s for 60-240 min in total with an image resolution of 1024 × 1024 pixel and a scan speed of 400 Hz. Transfection of the immune cells for transiently expressed proteins was accomplished with lipofectamin 2000 (Thermo Fisher Scientific) according to the manufacturer’s instructions. Live-cell analysis of heterologous expressed NFATc2-GFP and STIM1-eYFP/Orai1-eCFP localization was performed using the CLSM mentioned above. The microscopy settings were as follows - an image resolution of 1024 × 1024 pixel, scan speed of 200 Hz, time interval of 60 s for 30 – 100 min in total.

### FRET Analysis

FRET experiments with Jurkat cells, transiently expressing STIM1-eYFP/Orai1-eCFP, were examined with a Leica TCS SP5 II confocal microscope. Filter were set with CFP (458 Ex/460-490 Em), YFP (514 Ex/530-550 Em), and FRETraw (458 Ex/530-550 Em). Live-cell images were obtained every 30 s at room temperature with a 100 × 1.44 oil UV objective (HCX PL APO CS) for a time period of 30 min. Three-channel corrected FRET was calculated based on the following equation: FRETc = Fraw − Fd/Dd·FCFP − Fa/Da·FYFP, where FRETc represents the corrected total amount of energy transfer; Fd/Dd is the measured bleed-through of eCFP via YFP filter (0.473); Fa/Da represents measured bleed through of YFP through CFP filter (0.049). To reduce variations caused by differences in expression levels of CFP, the FRETc values were normalized to value of donor fluorescence (FCFP). To minimize the effect of variations of YFP expression levels on FCFP-normalized FRET signals (FRETN) and to show the relative changes as compared with resting levels, figures are shown as ΔFRETN/FRETNrest.

### Immunofluorescence staining

Jurkat cells were fixed on coated glass coverslips at 15, 30, 45, 60 or 90 min post treatment using 4% paraformaldehyde and next stained with primary antibodies for STIM1 (#PA1-46217, Thermo Fisher Scientifc), Orai1 (#NBP1-75523, Novus Biologicals, Waltham, MA, USA) and NFATc2 (#MA1-025, Thermo Fisher Scientific). Antibodies were applied at a 1:200 dilution in PBS and coverslips were shaken over night at 4°C. Next, cells were incubated with anti-rabbit IgG Alexa488 secondary antibody (# A32731, Thermo Fisher Scientifc), anti-mouse IgG Alexa488 secondary antibody (# A32723, Thermo Fisher Scientifc) or anti-mouse IgG Alexa647 secondary antibody (# A32728, Thermo Fisher Scientifc).

### Statistics

Data are expressed as mean ± standard deviation (SD) or standard error (SE) of ≥ three independent experiments; number of biological replicates (n) or independent experiments (N) are given in the text. Significance was estimated with Student’s t-test. P values <0.05 (*), < 0.01 (**) and < 0.001 (***) are indicated in the figures.

## Supplementary Materials

Figure S1. Radiation-induced Ca^2+^_cyt_ oscillations and nuclear NFAT translocation are triggered by X-ray and abolished after inhibition of Ca^2+^ influx.

Figure S2. In resting Jurkat cells STIM1 and Orai1 are located in the ER and PM respectively

## Acknowledgments

This work was supported in part by the German Research Foundation (DFG: Graduate school 1657) and by the German Federal Ministry of Education and Research (BMBF, grants no. 02NUK050A, 02NUK050C and 02NUK050D, GREWISalpha). We thank Donald Gill (Pennsylvania State University) and Ralph Kehlenbach (University of Göttingen) for providing STIM1-eYFP/Orai1-eCFP and NFAT-GFP plasmids, respectively. Special thanks to Christine Gibhardt (University of Göttingen) for helpful suggestions.

## Author contributions

Experiments, DT, TSp, SF, TS, SH and BJ; data analysis, DT, TS, GT; conceptualization, CF, FR, BR, AM, and GT; writing original draft preparation, DT, SF, BR, AM, GT; writing review and editing, FR, CF, GT; funding acquisition, CF, FR, GT.

