## Supplementary material for "X-ray Irradiation activates immune response in human T-lymphocytes by eliciting a Ca^2+^ signaling cascade": Suplementary figures 1 and 2

### Supplementary Materials

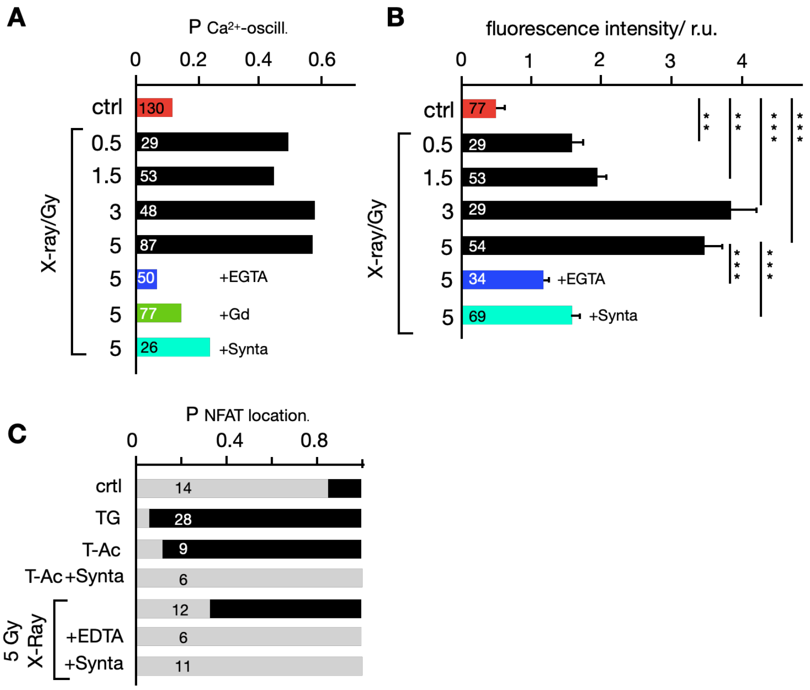


**Fig. S1: Radiation-induced $Ca^{2+}_{cyt}$ oscillations and nuclear NFAT translocation are triggered by X-ray and abolished after inhibition of $Ca^{2+}$ influx.**
(A) Probability for detecting $Ca^{2+}_{cyt}$ oscillations (P $Ca^{2+}$-oscill.) in Jurkat cells as in Fig. 1B. Cells were either non-irradiated (ctrl) or exposed to X-ray doses between 0.5 and 5 Gy in the absence (black bars) or presence of 5 mM EGTA (blue bar), 5 µM $Gd^{3+}$ (green bar) or 10 µM CRAC channel inhibitor Synta (turquoise bar). For analysis, only cells, which started oscillating ≥10 min after X-ray exposure were considered. (B) Maximal fluorescence intensity ± SD in non-irradiated cells (ctrl) and cells exposed to X-ray doses between 0.5 and 5 Gy in absence or presence of 5 mM EGTA or 10 µM Synta. The difference between treatments has a medium to high significance (** P< 0.01; *** P<0.001). (C) Probability of detecting NFAT in cytosol (gray bar) or in nucleus (black bar) as in Fig. 4A in untreated control cells (crtl), cells treated with 2 µM thapsigargin (TG) or with 25 µL/mL activator (T-Ac) without and with 10 µM Synta. Other cells were exposed to 5 Gy X-ray without or with 5 mM EGTA or 10 µM Synta. Numbers in A-C indicate the number of cells analyzed.

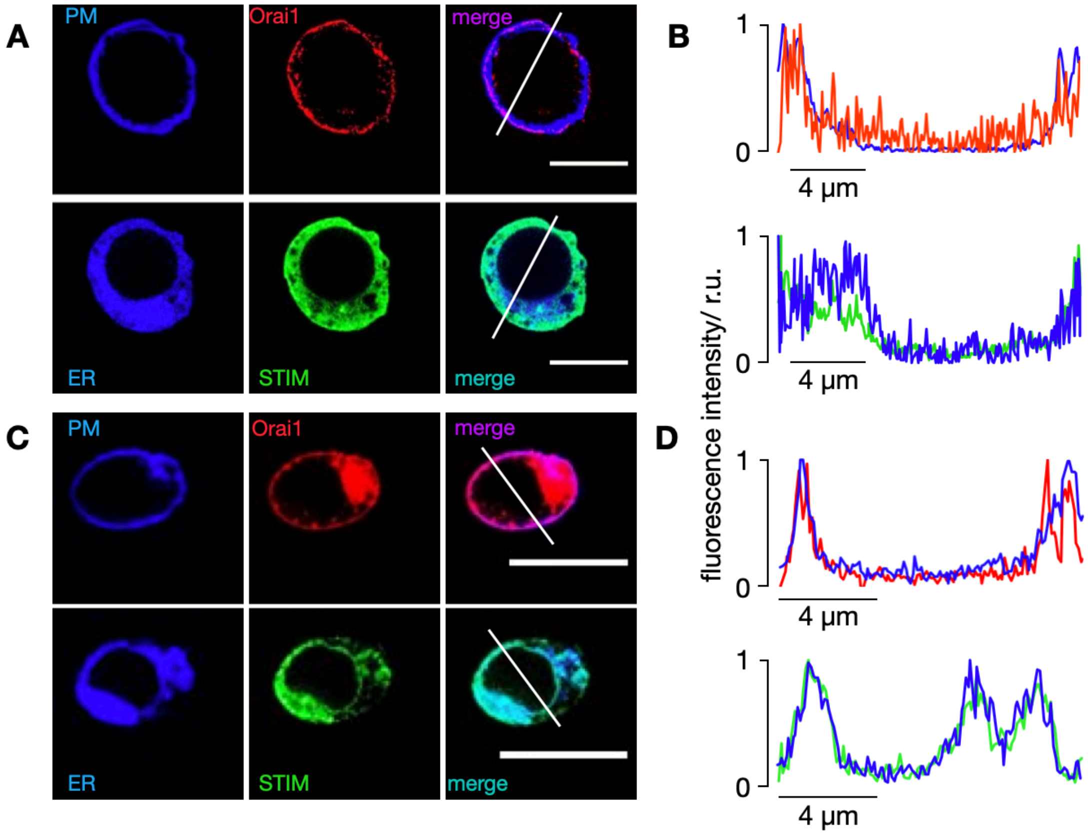


**Figure S2. In resting Jurkat cells STIM1 and Orai1 are located in the ER and PM respcetively**
Confocal images of cellular distribution of endogenous (A, lower row) STIM1 and (A, upper row) Orai1. Immunofluorescence was obtained by staining fixed (A) Jurkat cells and (C) PBMCs with Alx488 or Alx647 secondary antibody (central column). Additionally, the plasma membrane was visualized with the plasma membrane tracker CellMaskOrange and the ER with ER-tracker red (left column). An overlay of both channels is shown in right column. (B, D) Line Plots for each marker were taken in positions report in merged images. Fluorescence intensity of either (B) Orai1/PM or (D) STIM1/ER were normalized to the highest value of each signal; the colors of line plots correspond to those in images. All scale bars10 µm.
